# Theory of reinforcement learning and motivation in the basal ganglia

**DOI:** 10.1101/174524

**Authors:** Rafal Bogacz

**Affiliations:** MRC Brain Network Dynamics Unit, Nuffield Department of Clinical Neurosciences, University of Oxford, John Radcliffe Hospital, Oxford, OX3 9DU, UK

## Abstract

This paper proposes how the neural circuits in vertebrates select actions on the basis of past experience and the current motivational state. According to the presented theory, the basal ganglia evaluate the utility of considered actions by combining the positive consequences (e.g. nutrition) scaled by the motivational state (e.g. hunger) with the negative consequences (e.g. effort). The theory suggests how the basal ganglia compute utility by combining the positive and negative consequences encoded in the synaptic weights of striatal Go and No-Go neurons, and the motivational state carried by neuromodulators including dopamine. Furthermore, the theory suggests how the striatal neurons to learn separately about consequences of actions, and how the dopaminergic neurons themselves learn what level of activity they need to produce to optimize behaviour. The theory accounts for the effects of dopaminergic modulation on behaviour, patterns of synaptic plasticity in striatum, and responses of dopaminergic neurons in diverse situations.

## Introduction

In order to survive, animals need to select the most appropriate behaviour in a given situation. An important role in this process of action selection is played in all vertebrates by a set of subcortical structures called the basal ganglia (Redgrave et al., 1999). The information processing in the basal ganglia is very strongly modulated by dopamine. The basal ganglia are critically involved both in the process of selecting actions, and in learning which actions are worth making in a given context, as demonstrated by impairments of both functions in Parkinson’s disease. Death of dopaminergic neurons in Parkinson’s disease leads to problems with movements (Blandini et al., 2000) as well as difficulties in learning from feedback (Knowlton et al., 1996).

The basal ganglia is organized into two main pathways shown schematically in green and red in Figure 1. The Go or direct pathway is related to the initiation of movements, while the No-Go or indirect pathway is thought to be related to the inhibition of movements (Kravitz et al., 2010). These two pathways originate from two separate populations of striatal neurons expressing different dopaminergic receptors (Smith et al., 1998). The striatal Go neurons express D1 receptors and are excited by dopamine, while the striatal No-Go neurons express D2 receptors and are inhibited by dopamine (Surmeier et al., 2007). Thus dopamine changes the balance between the two pathways and promotes action initiation over inhibition.

**Figure 1:**
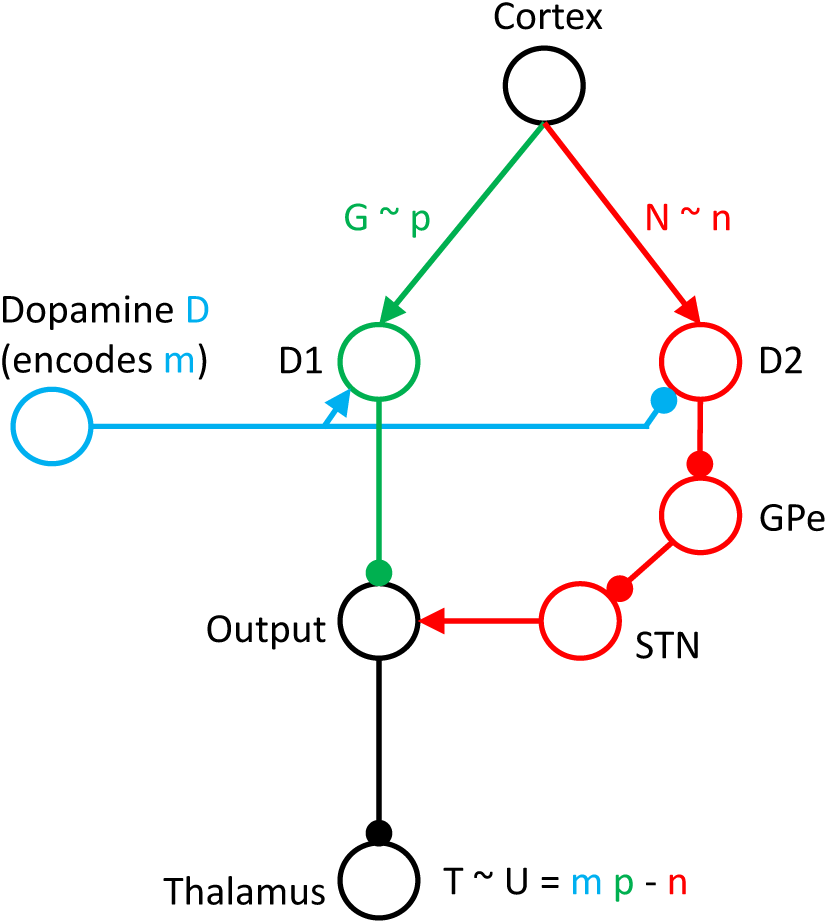
The organization of the basal ganglia. Circles denote neural populations in the areas indicated by labels next to them, where D1 and D2 corresponds to stiatal neurons expressing D1 and D2 receptors respectively, STN stands for the subthalamic nucleus, GPe for the external segment of globus pallidus, and Output for the output nuclei of the basal ganglia, i.e. internal segment of globus pallidus and substantia nigra pars reticulata. Arrows and lines ending with circles denote excitatory and inhibitory connections respectively.

The competition between Go and No-Go pathway during action selection and its dopaminergic modulation have been described by many computational models (e.g. Gurney et al., 2001; Humphries et al., 2012). In particular, the Opponent Actor Learning (OpAL) model suggested that the Go and No-Go neurons encode the positive and negative consequences of actions respectively, and they may bias the choice of action to a different extent depending on the level of dopamine, which encodes the current motivational state (Collins and Frank, 2014). On the other hand, the function of dopamine has been conceptualized in the incentive salience theories (Berridge and Robinson, 1998; Zhang et al., 2009). They propose that the learned values of stimuli are distinct from incentive values that determine choices, as the incentive values also depend on the current physiological state, that is encoded in dopaminergic activity. The first part of the Results aims at bringing together the models of direct and indirect pathways with the incentive salience theory in a single simple framework. It proposes that the computations of the basal ganglia can be formalized as evaluating a utility function, in which payoffs and costs (encoded by Go and No-Go neurons) are scaled differently depending on the motivational state (encoded by dopaminergic neurons).

Much research has focussed on how the synapses of Go and No-Go neurons are modified by experience. It has been observed that bursts of activity of dopaminergic neurons encode reward prediction error defined as the difference between reward obtained and expected (Schultz et al., 1997; Eshel et al., 2016). Such dopaminergic activity has been shown to produce distinct changes in the synaptic weights of Go and No-Go neurons (Shen et al., 2008). The process of learning in synapses of Go and No-Go neurons process has been described by several computational models (Frank et al., 2004; Hong and Hikosaka, 2011; Gurney et al., 2015; Yttri and Dudman, 2016). Among them, the OpAL model provided simple and analytically tractable rules describing the changes in weights of Go and No-Go neurons as a function of reward prediction error (Collins and Frank, 2014). However, it has not been shown if the weights of Go and No-Go neurons selective for an action can converge over trials to values proportional to its payoff and cost. In the second part of the Results, it is demonstrated that recently proposed learning rules (Mikhael and Bogacz, 2016) allow the Go and No-Go neurons to estimate both payoffs and costs associated with a given action.

Dopaminergic neurons have also been reported to respond in other situations, such as after presentation of salient stimuli (Redgrave and Gurney, 2006), novel stimuli (Schultz, 1998), aversive stimuli (Matsumoto and Hikosaka, 2009), reward uncertainty (Fiorillo et al., 2003), and during movements (Howe and Dombeck, 2016). It is challenging for the classical reinforcement learning theory to account for all aspects of activity of dopaminergic neurons (Syed et al., 2016). In the third part of the Results section, it is argued that these diverse responses arise because the dopaminergic neurons themselves learn to responds in states in which acting gives higher reward than not acting.

The theory proposed in the paper accounts for the effects of dopamine depletion on behaviour, the dopaminergic modulation of synaptic plasticity, and for the responses of dopaminergic neurons in diverse situations.

## Results

### Utility of actions

Let us consider how a system controlling behaviour of an animal or a human needs to operate to maximize their chances for survival. Any action may have some positive and negative consequences. For example, eating an apple has positive consequences, as it gives nutrients and water. But it may also have a cost, as in order to get the apple one may need to climb a tree which costs metabolic energy, and on the way one may be attacked by a predator. While evaluating the utility of an action, the brain needs to combine the positive and negative consequences, which we denote by *p* and *n*. However, the value of positive consequences also depends on the current level of motivation, e.g. hunger. The nutrients in the apple are only valuable if one’s food reserves are low. Let us denote the motivation by *m*, and consider the following simple utility function:

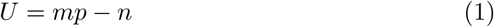

In this utility equation motivation scales the positive consequences, as in one of the incentive salience models (Zhang et al., 2009). However, motivation does not scale the negative consequences because being attacked by a predator is equally bad no matter if one is hungry or not.

Let us now consider how the brain should select actions. The brain should evaluate the utility of available actions, and choose the one with the highest utility. But this best action should only be executed if its utility is higher than the utility of doing nothing. Otherwise one should not take any actions. For example, consider a lion who just ate a big antelope. She is not hungry, i.e. *m* = 0, so according to Equation 1, any action involving some cost or effort will have negative utility, thus the lion should just relax.

We propose that the utility of actions is evaluated in the basal ganglia, and that the architecture of this circuit exactly matches that necessary to compute the utility function, in which the motivation differentially scales the positive and negative consequences.

Following the OpAL model (Collins and Frank, 2014), we assume that the positive consequences of choosing a particular action in a particular state are encoded in the synaptic weights of connections from the cortical neurons selective for the state and the striatal Go neurons selective for the action. These weights are denoted by *G* in Figure 1. We propose that after learning, the weight *G* is proportional to *p*, i.e. *G* = *cp* where *c* is a constant, which we denote for brevity by *G ∼ p*. In the model, the negative consequences are encoded in the synaptic connections of striatal No-Go neurons. We denote their weights by *N*, and assume that after learning they are proportional to the negative consequences *n*. The motivational state *m* is encoded in the model by the level of dopaminergic activity, which we denote by *D*. It is plausible to assume that the dopaminergic activity is modulated by motivation, because receptors regulating food intake are expressed in areas including dopaminergic neurons, and satiety signals inhibit dopamine release, while feeding promotors enhance dopamine signalling (Phillips et al., 2007). Moreover, thirst increases levels of dopamine in the brain (Zabik et al., 1993), and the responses of dopaminergic neurons depend on hunger and thirst (Papageorgiou et al., 2016).

As dopaminergic neurons modulate the Go and No-Go neurons in opposite ways, dopamine can control to what extent the positive and negative consequences affect the basal ganglia output. We will now demonstrate that, thanks to this modulation, the activity in the thalamus can be proportional to the utility function. Let us start by assuming that the weights of striatal neurons have already been learned and are proportional to the positive and negative consequences (we will show how this learning can occur in the next subsection). Let us now think how the thalamic activity, which we denote by *T*, depends on the cortico-striatal weights and dopaminergic modulation. This relationship is surely complex, but let us write a simple equation that just captures the signs of the dependencies:

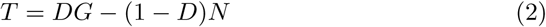

In the above equation the first term *DG* correspond to the input from the striatal Go neurons. This term is positive, because the projection from striatal Go neurons to the thalamus involves double inhibitory connection (see Figure 1) so it overall has excitatory effect. The activity of the Go neurons depends on synaptic weights *G*. We assume that their gain is modulated by the dopaminergic input *D*, based on the observation of an increased slope of the firing-input relationship in the presence of dopamine (Thurley et al., 2008). The second term −(1 − *D*)*N* corresponds to the input from the striatal No-Go neurons. It has a negative sign because the projection form the No-Go neurons to the thalamus includes three inhibitory connections. The activity of the striatal No-Go neurons depends on their synaptic weights *N*, and we assume that their gain is reduced by dopamine, so the synaptic input is scaled by (1 − *D*). In Equation 2, we assume that *D* ϵ [0, 1], and the value of *D* = 0.5 corresponds to a baseline level of dopamine for which both striatal populations equally affect the thalamic activity.

Let us now show that the thalamic activity defined in Equation 2 is proportional to the utility function of Equation 1. Substituting *G ∼ p*, *N ∼ n* into Equation 2 and rearranging terms we obtain:

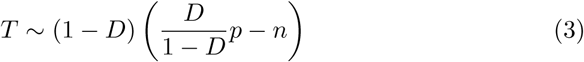

Comparing Equations 1 and 3, we see that *T* is proportional to *U* when motivation is encoded in the following function of dopaminergic activity:

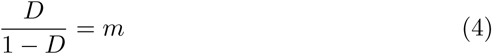

Rearranging terms, we see how the dopaminergic activity should depend on the level of motivation:

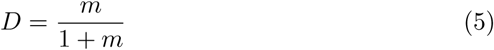

In summary, when the striatal weights encode the consequences and the dopamine level is described by Equation 5, then the thalamic activity is proportional to the utility. The cortico-basal-ganglia-thalamic circuit includes neurons selective for different actions, and the activity of thalamic neurons selective for specific actions is determined on the basis of their individual payoffs and costs and the common dopamine level. As the proportionality coefficient (1 − *D*) in Equation 3 is the same for all actions, the most active neurons in the thalamus are the ones selective for the action with the highest utility, so this action may be chosen through competition. Furthermore, if we assume that actions are only selected when thalamic activity is above a threshold, then no action will be selected if all actions have insufficient utility.

In order to evaluate the utility in Equation 1, in which the motivation differently scales positive and negative consequences, the payoffs and costs need to be stored separately. If a single synaptic weight was already encoding the difference *p − n*, then it would not be possible to scale just *p* by the motivation. This shows that any neural system computing the utility of the form considered here needs to include separate sets of synapses to encode *p* and *n*. This may be a fundamental reason why the basal ganglia include both Go and No-Go pathways.

It may seem surprising that dopaminergic neurons modulate both Go and No-Go neurons, although the motivation term only scales *p* in the definition of the utility function of Equation 1. To understand why such double modulation is necessary, let us consider a situation in which an animal is extremely hungry and eating something immediately is necessary to prevent starvation. In such a situation the animal must take a chance to execute the action irrespectively of the cost, as inaction would result in certain death. As above, let us assume that actions are only selected when the thalamic activity is above a fixed threshold. If the dopaminergic neurons only modulated the Go neurons, then the thalamic activity would be equal to the utility *U*. To ensure that *U* is above threshold for any *n*, the motivation *m* would have to be infinite (to guarantee that *mp* always outweighs *n* - see Section 2 of Supplemental Information). By contrast, to completely ignore the effects of negative consequences on thalamic activity defined in Equation 2, it is sufficient to set the dopamine level to *D* = 1. Thus, thanks to dopamine modulating both the Go and No-Go neurons, the required range of dopaminergic modulation is reduced from an unrealistically wide range of [0, *∞*] to a bounded range of [0, 1].

For simplicity, we considered the utility of Equation 1, in which the motivation multiplies positive consequences and does not scale negative consequences. However, Section 1 of Supplemental Information shows that the basal ganglia circuit could also evaluate a general class of utility functions in which the positive and negative consequences are differentially scaled by motivation. In the Results section we consider a simple case in which consequences and motivation have a single dimension, and in the Discussion we will come back to extending the theory to the case of multiple dimensions of *p* (e.g. food and water), *m* (e.g. hunger and thirst) and *n* (e.g. effort and pain).

Let us consider how the proposed theory relates to the effects of dopamine depletion on behaviour. In a classic experiment illustrated in Figure 2A, rats were given a choice between pressing a lever 5 times in order to obtain a tasty reward, and freely available lab chow (Salamone et al., 1991). Normal animals were willing to work for tasty food, but after dopamine depletion they were not willing to make an effort and preferred a less valuable but free option. A mechanical explanation for this surprising effect has been provided by Collins and Frank (2014). The theory proposed in this paper accounts for it in a conceptually similar but slightly simpler way, which is summarized below (simulations are described in the next subsection, and differences to the account of OpAL model in Discussion).

**Figure 2:**
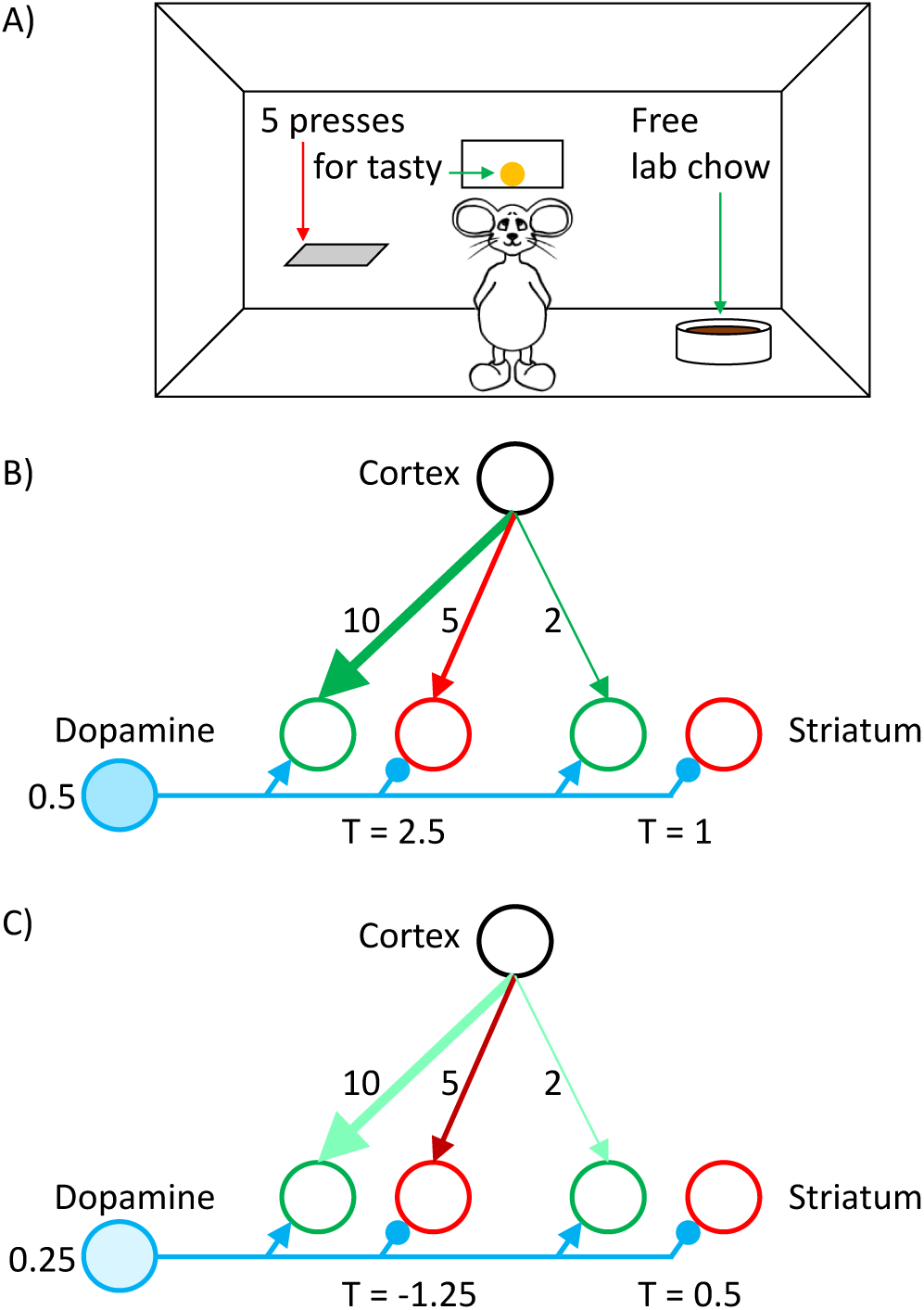
Critical role of dopamine in motivation. A) Schematic illustration of experimental paradigm − see text. B) Computation of utility in dopamine intact state. Green and red circles on the left denote striatal Go and No-Go neurons associated with pressing the lever, while the green and red circles on the right denote the neurons associated with approaching free food. The strength of the synaptic connections is denoted by the thickness of the arrows, and labels. These weights were set to sample values allowing easy explanation of the observed effect. The blue circle represents a population of dopaminergic neurons, and its shading indicates the level of activity. C) Computation of utility in dopamine depleted state. Notation as in panel B, but additionally the light green colour of the connection of the Go neurons indicates that their gain has been reduced, while the dark red color of the connections of the No-Go neurons symbolizes an increased gain.

Figure 2B illustrates how the model can account for the behaviour when the dopamine level has a normal baseline value. In the figure, the strength of the cortico-striatal connections is denoted by the labels and the thickness of arrows. Pressing the lever gives a high payoff, so the weights of Go neurons selective for this action are strong, but it also has a substantial cost, so the No-Go weights are also present. On the other hand, the free food is not particularly tasty so the Go weights are weak, and there is no cost, so the No-Go weight is *N* = 0 (represented by the lack of connection to the rightmost striatal population in Figure 2B). When the dopamine level is at baseline, the positive and negative consequences are weighted equally, so the thalamic neurons selective for pressing the lever have overall higher activity (computed from Equation 2), so this action is more likely to be chosen.

By contrast, Figure 2C shows that when the dopamine level is reduced, costs are weighted more than payoffs, and the utility of pressing the lever becomes negative. As approaching free food does not have any cost, the activity of thalamic neurons selective for this option is now higher, and this action is more likely to be chosen.

The unwillingness to make an effort for reward in dopamine depleted state has also been observed in other paradigms: During a choice in a T-maze, dopamine depleted animals were less likely to go to an arm with more pellets behind the barrier, but rather chose the arm with easily accessible but fewer pellets (Salamone et al., 2016). Parkinson’s patients were not willing to exert as much physical effort by squeezing a handle in order to obtain reward as healthy controls, especially if they were off medications (Chong et al., 2015). These effects can be explained in an analogous way (Collins and Frank, 2014) by assuming that in the dopamine depleted state the effort of crossing the barrier or squeezing a handle is weighted more, resulting in lower activity of thalamic neurons selective for this option. According to the theory proposed here, reducing the dopamine level reduces the utility of actions involving costs, and thus changes preferences.

### Learning the consequences of actions

This subsection shows that previously proposed plasticity rules (Mikhael and Bogacz, 2016) allow the Go and No-Go neurons to learn the positive and negative consequences of actions.

It has been proposed that striatal neurons learn on the basis of the reward prediction error signal encoded in bursts of firing of dopaminergic neurons (Schultz et al., 1997; Eshel et al., 2016). However, if the dopaminergic neurons carry both the motivation and teaching signals, the striatal neurons need to have a way to distinguish what signal is encoded at the moment and react appropriately, i.e. change their gain according to the motivation signal, and change the synaptic weights according to the teaching signal. Although it has been suggested that the motivation is encoded in the tonic dopamine level, while the reward prediction error in the phasic changes in activity (Niv, 2007), this distinction has been recently questioned (Howe and Dombeck, 2016; Hamid et al., 2016), and in the next subsection we will propose that motivational signal *m* may also change on a fast time scale. The information on whether the dopaminergic neurons encode motivation or teaching signals in a given moment may be provided by other means. For example, it has been suggested that the cholinergic neurons provide a permissive input, which enables plasticity of the cortico-striatal synapses (Deffains and Bergman, 2015), so their level of activity may inform what the dopaminergic neurons encode at the moment. For simplicity, from now on we will refer to the dopaminergic activity encoding *m* and reward prediction error as the dopaminergic motivation and teaching signals respectively, and we will come back to possible ways the striatal neurons can distinguish between them in the Discussion.

To learn separately about positive and negative consequences of actions, the striatial neurons can take advantage of the fact that these consequences typically occur in different moments of time. Let us consider a situation in which an animal performs an action involving a cost *n* in order to obtain a payoff *p* (e.g. pressing a lever in order to obtain tasty food, as in Figure 2A). The changes in the instantaneous reinforcement *r* during the course of this action are schematically illustrated in Figure 3A (in the lever pressing example, the reinforcement is negative while pressing the lever due to the effort and then positive, when the payoff is obtained). The prediction error would also take negative and then positive values (a negative prediction error is thought to be encoded in a decrease of firing of dopaminergic neurons below the baseline (Schultz et al., 1997)).

**Figure 3:**
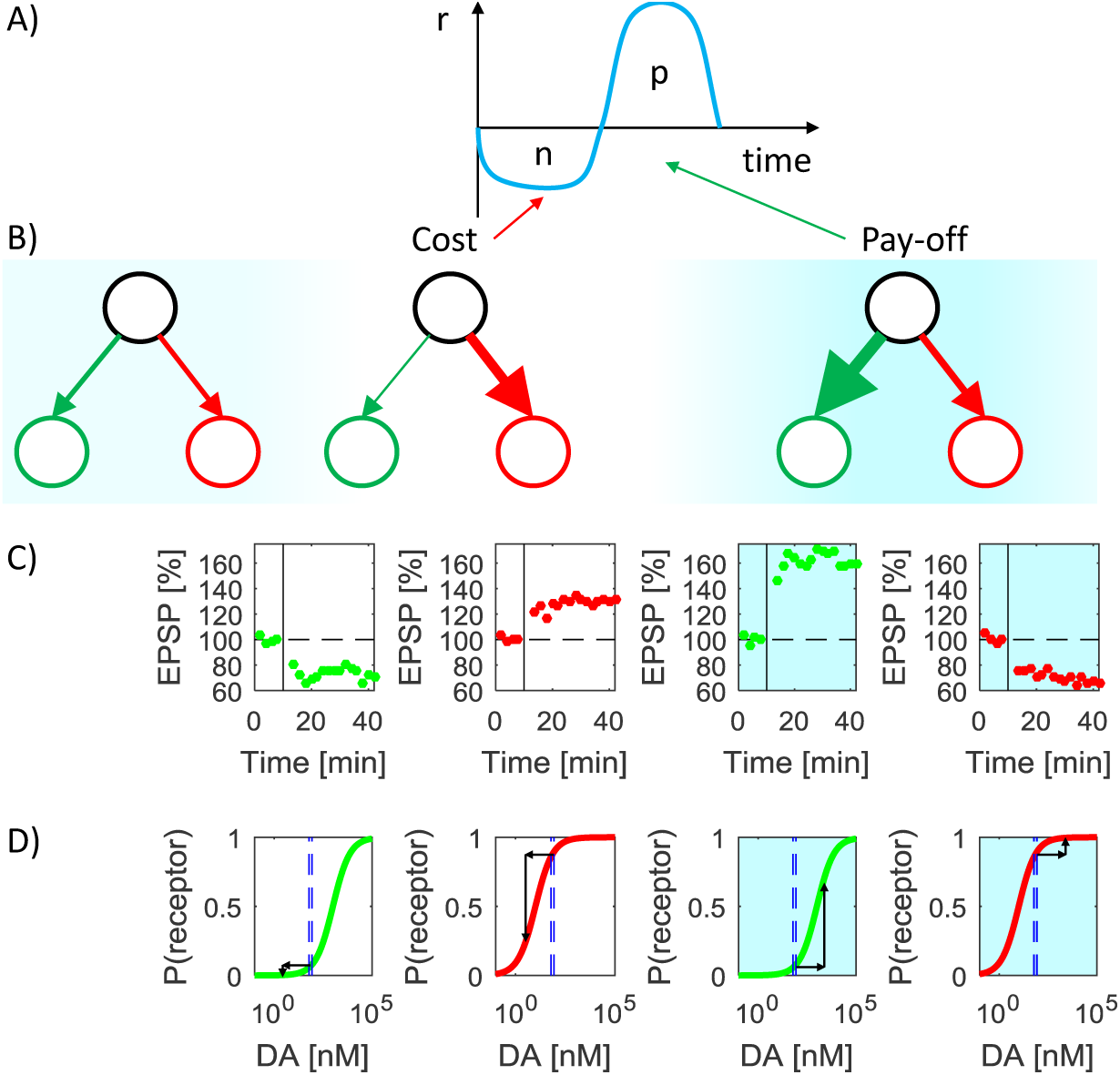
Learning the positive and the negative consequences of actions. A) Instantaneous reinforcement *r* when an action with effort *n* is selected to obtain payoff *p*. B) Cortico-striatal weights before the action, after performing the action, and after obtaining the payoff. Red and green circles correspond to striatal Go and No-Go neurons, and the thickness of the lines indicates the strength of synaptic connections. The intensity of the blue background indicates the dopaminergic teaching signal at different moments of time. C) The average excitatory post-synaptic potential (EPSP) in striatal neurons produced by cortical stimulation as a function of time in the experiment by Shen et al. (2008). The vertical black lines indicate the time when the synaptic plasticity was induced by successive stimulation of cortical and striatal neurons. The amplitude of EPSPs is normalized to the baseline before the stimulation indicated by horizontal dashed lines. The green and red dots indicate the EPSPs of Go and No-Go neurons respectively. Displays with white background show the data from experiments with rat models of Parkinson’s disease, while the displays with blue background show the data from experiments in the presence of corresponding dopamine receptor agonists. The four displays re-plot the data from Figures 3E, 3B, 3F and 1H in the paper by Shen et al. (2008). D) Changes in dopamine receptor occupancy. The green and red curves show the probabilities of D1 and D2 receptor occupancies in a biophysical model of Dreyer et al. (2010). The two dashed blue lines in each panel indicate the levels of dopamine in dorsal (60 nM) and ventral (85 nM) striatum estimated on the basis of spontaneous firing of dopaminergic neurons using the biophysical model (Dodson et al., 2016). Displays with white and blue backgrounds illustrate changes in receptor occupancy when the level of dopamine is reduced or increased respectively.

Let us first provide an intuition for how the plasticity rules operate. Figure 3B sketches the changes in the synaptic weights. The leftmost display shows the initial weights. While making an effort to perform an action, the reward prediction error is negative. Similarly as in previous models (Frank et al., 2004; Collins and Frank, 2014), we assume that the negative prediction error results in an increase in *N* (compare the red arrows in the middle and the left displays in Figure 3B). This allows weights *N* to encode negative consequences *n*. Following payoff, the prediction error is positive, and in the model *G* increases, allowing weights *G* to encode *p*. So if an action involves both positive and negative consequences, both weights are increased (compare the right and the left displays in Figure 3B). To prevent the weights from increasing to infinity, we also assume that the weights decay (so weights that did not increase are made smaller in subsequent displays in Figure 3B).

Let us now formalize the above description of striatal plasticity. To understand why the plasticity rules need to have their particular form, it is helpful to first consider simpler hypothetical plasticity rules that are easier to analyse. Let us for a moment make a simplifying assumption that dopaminergic teaching signal encodes instantaneous reinforcement rather than reward prediction error (we will consider the prediction error in a moment). Under this assumption the following plasticity rules could learn the positive and negative consequences.

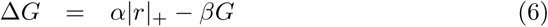

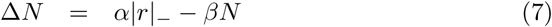

The update rule in each of the above equations consists of two terms. The first term is the change depending on the dopaminergic teaching signal scaled by a learning rate constant *α*. In the rule for Go neurons |*r*|_+_ denotes the reinforcement when *|r|* > 0 and 0 otherwise, so these neurons increase weights only if the reinforcement is positive. Conversely, in the rule for No-Go neurons *|r|*_-_ denotes the absolute value of the reinforcement when *r <* 0 and 0 otherwise, so these neurons only increase their weights when the reinforcement is negative. The second term in the update rules is a decay term, scaled by a decay rate constant *β*. This term is necessary to ensure that the weights stop growing when they are sufficiently high.

Let us now show that when the weights *G* and *N* are modified according to these rules in the scenario shown in Figure 3A, they converge to the values proportional to *p* and *n*. When an action has a cost *n* and payoff *p*, the weights are updated twice: with *r* = *−n* after making an action, and *r* = *p* after receiving the payoff. Thus the weight changes are approximately equal to:

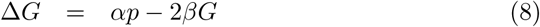

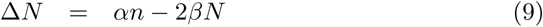

There is a factor of 2 before the decay term as the decay happens both during making an effort and receiving the payoff. Once the weights converge after performing the action multiple times, the change in weights must be Δ*G* = Δ*N* = 0. Setting the left hand side of the above equations to 0 and solving for *G* and *N*, we obtain the values to which the weights converge (which we indicate with a star):

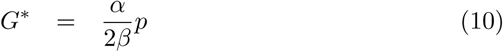

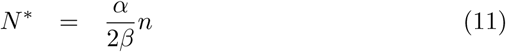

We see that the weights converge to the values proportional to the positive and negative consequences. At these values, the weight increases due to cost and payoff are exactly balanced by the decay, so the weights do not change from trial to trial anymore.

Let us now consider how the plasticity rules of Equations 6 and 7 need to be adjusted when we make a more realistic assumption that the dopaminergic teaching signal encodes the reward prediction error defined as the difference between reinforcement obtained *r* and that expected. Throughout the main text of the paper we assume that the expected reward for selecting a particular action is encoded in the difference of weights of Go and No-Go neurons selective for this action, i.e. *G − N*. Thus we can define the reward prediction error as:

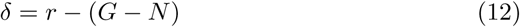

In Section 4 of Supplemental Information it is shown that the theory described in the main text also generalizes to the actor-critic framework (Doya, 1999) that assumes that the expected reward is computed by a separate group of striatal patch neurons.

When the dopaminergic teaching signal encodes reward prediction error, the changes in synaptic weights need to take the following form (Mikhael and Bogacz, 2016):

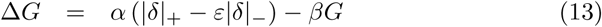

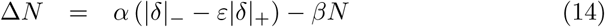

The rules include an additional term scaled by a constant 0 *< ε <* 1. Thus, the weights of Go neurons are increased when *δ >* 0, but also slightly decreased when *δ <* 0, and the constant *ε* controls the magnitude of this additional decrease. To illustrate the need for this decrease, Figure 4 shows reward prediction error in two cases which have the same payoff but differ in costs. It is very intuitive that the initial decrease is more pronounced for the action with higher cost (shown in purple), but this action also produces higher prediction error while receiving the payoff, despite the payoff being the same for the two actions. This happens because the action with the higher cost has a lower overall value so there is a larger difference between the payoff and the expected value. If the weights were modified by the rules without the additional term (or with *ε* = 0), then *G* would converge to a slightly higher value for the option with the higher cost, despite the same payoff for both actions. To ensure that the weight *G* only depends on payoff, it needs to also slightly decrease when *δ <* 0.

**Figure 4:**
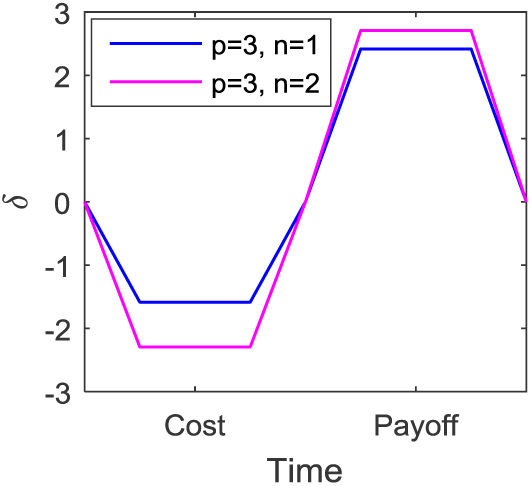
Changes in reward prediction error during execution of two actions which differ in cost *n* but have the same payoff *p* (see key). The figure shows the prediction errors <u>o</u>f Equation 12 produced after convergence by a model with *α* = *β* and 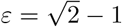. The values of expected reward during cost and payoff were computed from Equation 21.

In the Materials and Methods section it is shown analytically that for an action with payoff *p* and cost *n*, the weights *G* and *N* converge to values proportional to *p* and *n* respectively, when the parameters of the learning rule (*α*, *β*, *ε*) satisfy a particular relationship (Equation 25). This property is illustrated in the simulations shown in Figure 5. In each column simulations were performed with the same payoff, and weights *G* converge to the same values within a column. Analogously, in each row the simulations were performed with the same *n*, resulting in the same values of *N* within a row.

**Figure 5:**
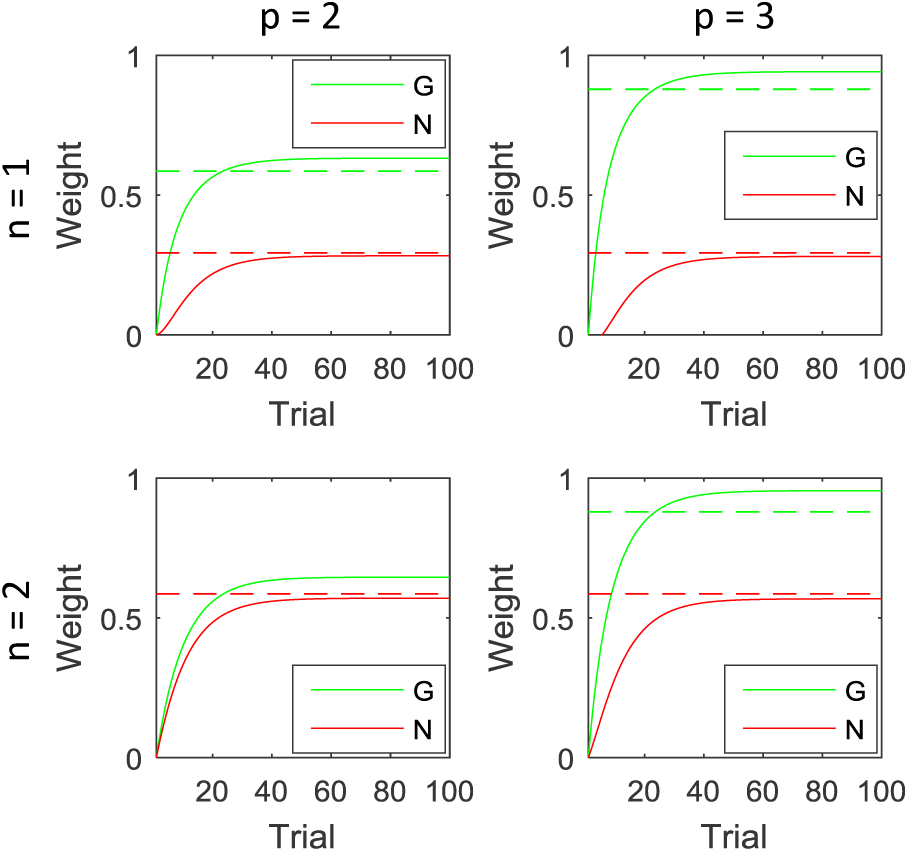
Changes in the weights *G* of Go neurons and *N* of No-Go neurons over the course of simulations. Each display shows the results of a simulation with different *p* and *n*: in the left displays *p* = 2, while in the right displays *p* = 3; in the top displays *n* = 1, while in the bottom displays *n* = 2. The simulations were run with parameters *α* = *β* = 0.05 and 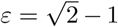. The dashed lines indicate the positions of the fixed points computed from Equation 26 for *G* or an analogous equation for *N*.

In Section 3 of Supplemental Information, we show that the computation of utility is robust to variations in parameters. Namely, even if the parameters do not precisely satisfy the condition of Equation 25, the weights converge to linear combinations of *p* and *n*, which still allows the circuit to evaluate the utility of actions (but with a less orthogonal coding of *p* and *n* in *G* and *N*).

Let us now consider how the weight changes illustrated in Figure 3B and formalized in Equations 13 and 14 relate to known data on synaptic plasticity in the striatum. The direction of changes in *G* and *N* depending on the sign of *δ* are consistent with the changes of synaptic weights of Go and No-Go neurons observed at different dopamine concentrations. Figure 3C shows experimentally observed changes in synaptic strengths when the level of dopamine is low (displays with white background) and in the presence of agonists (blue background) (Shen et al., 2008). Note that the directions of change match those in the corresponding displays above, in Figure 3B.

These directions of changes in striatal weights are also consistent with other models of the basal ganglia (Frank et al., 2004; Collins and Frank, 2014), but the unique prediction of the rules of Equations 13 and 14 is that the increase in dopaminergic teaching signal should mainly affect changes in *G*, while the decrease in dopamine should primarily affect *N*. Thus, the dopamine receptors on the Go and No-Go neurons should be most sensitive to increases and decreases in dopamine level respectively. This matches with the properties of these receptors. The D2 receptors on No-Go neurons have a higher affinity and therefore are sensitive to low levels of dopamine compared to D1 receptors on Go neurons (Richfield et al., 1989). This property is illustrated in Figure 3D where the green and red curves show the probabilities of D1 and D2 receptors being occupied as a function of dopamine concentration. The blue dashed lines indicate the levels of dopamine in the striatum predicted to result from spontaneous firing of dopaminergic neurons (Dodson et al., 2016). At these levels most D1 receptors are deactivated. Thus the D1 receptor activation will change when the dopamine goes up, but not when it goes down, as indicated by the black arrows. This is consistent with a higher effect of positive *δ* on weight changes of Go neurons in Equation 13. By contrast the D2 receptors are activated at baseline dopamine levels, so their activation is affected by the decreases in dopamine level but little by increases, in agreement with a higher effect of negative *δ* on No-Go neurons in Equation 14. In summary, the plasticity rules allowing learning positive and negative consequences are consistent with the observed plasticity and the receptor properties.

We now demonstrate that a model employing the proposed learning rules can quantitatively account for the effects of dopamine depletion on behaviour illustrated in Figure 2. Left displays in Figure 6 summarize experimental data (Salamone et al., 1991). Top-left display corresponds to a condition in which both tasty pellets and the lab chow were freely available. In this case, the animals preferred pellets irrespectively from dopamine level. The bottom-left panel corresponds to the condition in which the animal had to press a lever 5 times in order to obtain a pellet, and as mentioned before, after injections of a dopamine antagonist they started to prefer the lab chow.

**Figure 6:**
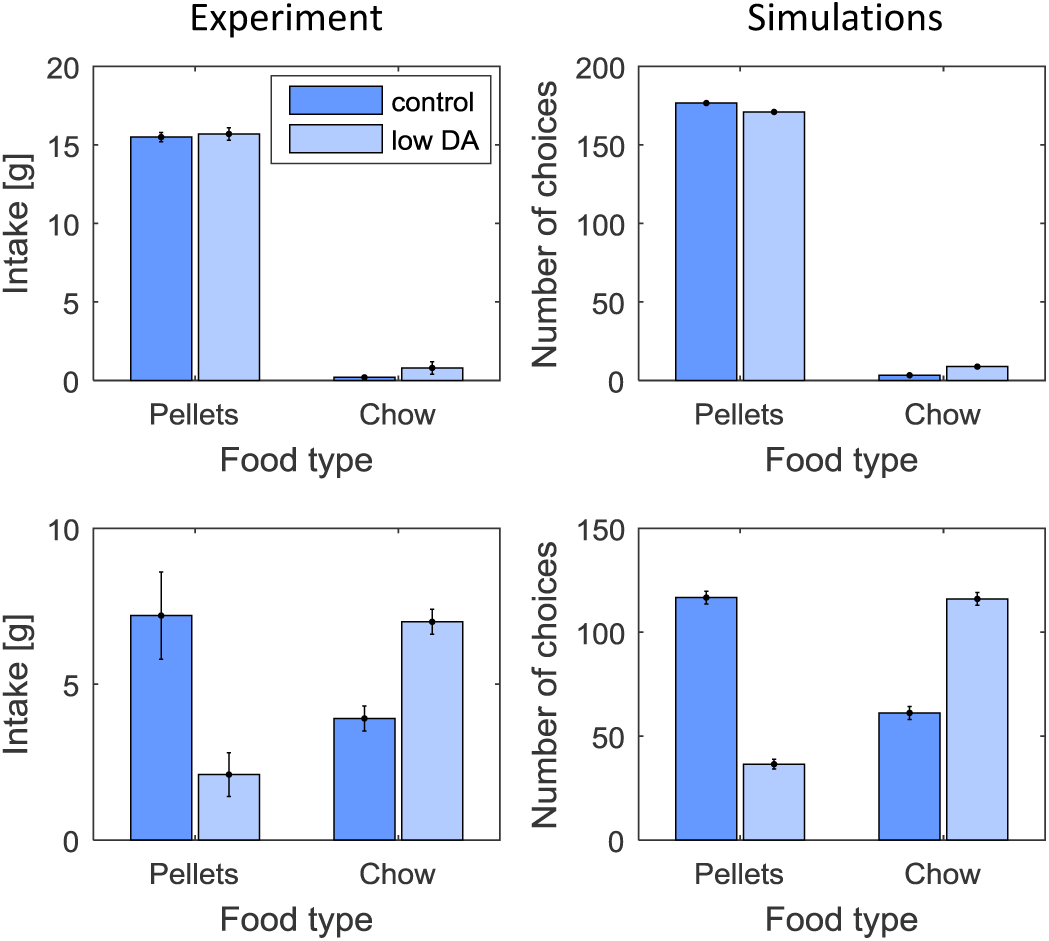
Frequency of choosing pellets and lab chow in dopamine intact (dark blue) and dopamine depleted (light blue) states. Top displays correspond to a condition with free pellets, while bottom displays to a condition where pressing a lever 5 times was required to obtain a pellet. Left displays re-plot experimental data. The values in the top-left and bottom-left displays were taken from Figures 1 and 4 respectively in the paper by Salamone et al. (1991). The right displays show the results of simulations. Error bars indicate the standard error of the mean.

As illustrated in Figure 2, the model assumes that on each trial the animal makes a choice between two actions: pressing a lever or approaching lab chow. Before the main experiments, the animals were trained to press lever to obtain reward and were exposed to the lab chow (Salamone et al., 1991). To parallel this in simulations, the model was first trained such that it experienced each action a number of times, received corresponding payoffs and costs, and updated its weights according to Equations 13 and 14. Then the model was tested with baseline and reduced dopaminergic motivation signal. As described in Materials and Methods, the parameters of the model were optimized to match experimentally observed behaviour. As shown in the right displays in Figure 6, the model was able to reproduce the observed pattern of behaviour. This illustrates model’s ability to capture both learning about payoffs and costs associated with individual actions and the effects of dopamine level on choice processes.

### Learning the motivation signal

This subsection proposes that the dopaminergic neurons themselves learn what motivation signal *m* they need to produce. As the level of dopamine controls the balance between the Go and No-Go pathways, increasing dopamine level raises the extent to which the motor system is “energized” and thus increases a general tendency to perform actions. If so,then to maximize reward, the dopaminergic neurons could adjust their motivation signal, and respond in states of the environment in which acting is generally useful, i.e. in which acting in general yields higher reward than not acting. The dependence of dopaminergic motivation signal on an extrinsic state of environment in addition to the intrinsic state is consistent with the observations that the tendency to search for food does not only depend on animal’s food reserves, but also on external circumstances like the presence of predators and food availability (Mrosovsky and Sherry, 1980). In this subsection we demonstrate that adjusting dopaminergic motivation signal on the basis of the state of the environment allows the basal ganglia to closer approximate the optimal policies, and may explain the diversity of situations in which the dopaminergic neurons have been observed to respond.

To see the benefit of adjusting the motivation signal according to the state of environment, let us consider an example of foraging. An animal that collects fruits from trees needs to make decisions which tree to approach. The utility of foraging on a tree clearly depends on the density of fruits on that tree. But it also depends on the amount of daylight, as it is difficult to collect fruits in the darkness. Assuming that no fruits can be collected when it is completely dark, the utility of foraging on a tree can be written as:

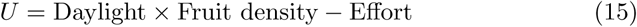

The above equation has the same form as the utility defined in Equation 1. Here the factor determining the benefit of foraging is the extrinsic state of daylight rather than the intrinsic state of hunger. This utility can be evaluated in the basal ganglia, if the dopaminergic neurons provided the information about the time of the day. Then the low level of dopaminergic motivation signal can prevent foraging during the night in an analogous way it prevents it when an animal is not hungry. Indeed, dopamine has been reported to be among other neuromodulators involved in wakefulness regulation (Wisor et al., 2001). Intrinsic factors such as hunger of course also affect the utility (and could be introduced into the above equation), so the dopaminergic motivation signal needs to depend on a combination of intrinsic and extrinsic factors. This example illustrates that it is particularly beneficial for the dopaminergic motivation signal to depend on the states of the environment that change a general structure of how reinforcements depend on actions.

The states in which acting is beneficial tend to be similar across animals, so much of the optimization of the motivation signal could have occurred on the evolutionary time-scale. Nevertheless, the dopaminergic motivation signal could be also further refined during animal’s lifetime to adjust it to particular circumstances faced by the animal. To identify the patterns of dopaminergic activity that optimize behaviour, and to illustrate that they can be tuned by experience, we simulated learning of response of dopaminergic neurons according to a classic model called Reinforce (Williams, 1992). The Reinforce model describes a learning rule that allows a neuron to produce the activity level that maximizes reward (Williams, 1992). Thus even if the tendency of dopaminergic neurons to respond in particular situations were a result of evolution, the simulations inform what activity the dopaminergic neurons are expected, if they were tuned over generations to maximize reward.

The Reinforce model (Williams, 1992) assumes that the neurons learn by trial and error what level of activity they need to produce to maximize reward. The model assumes that a dopaminergic neuron receives input from neurons encoding the current state or context (Figure 7A). The weights of synaptic connections from the state neurons determine the average activity of the dopaminergic neuron. But the neuron is stochastic and may produce higher activity on some trials and lower activity on others (Figure 7B). This noise allows the dopaminergic neuron to explore the effects of changes in its activity on reward: If the reward is higher than expected, the weights are modified to make the produced activity more likely (Figure 7C). If the reward were lower than expected, the weight changes would be opposite (see Materials and Methods for details).

**Figure 7:**
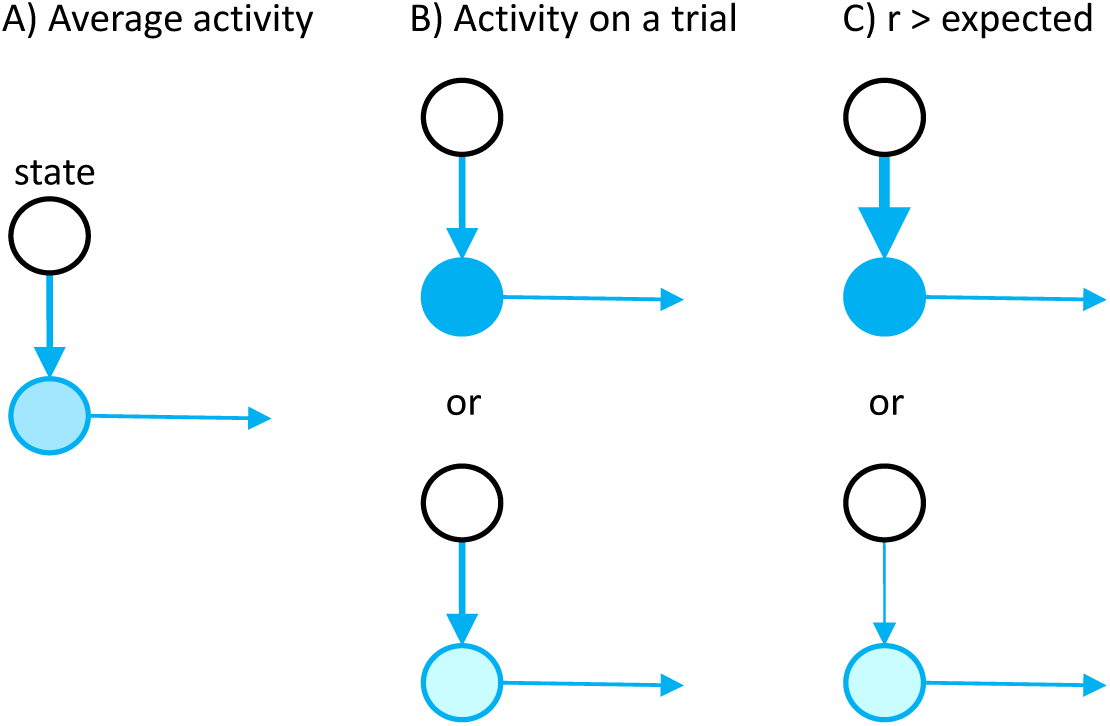
Learning in the Reinforce model. Blue circles denote a dopaminergic neuron, and their shading denotes the level of activity. Black circles denote a population of neurons providing information on the current state or context. The thickness of arrows indicates weights of synaptic connections. A) Average activity of the neuron. B) Despite the same synaptic weight, the neuron may produce higher (top display) or lower (bottom display) activity on individual trials. C) Changes in synaptic weights when reward is higher than expected. If the neuron produced higher activity *D* (top display), its weights are increased to make it more active in the future. If the neuron produced lower activity (bottom display), the weights are decreased.

Let us come back to the example of the influence of daylight on the utility of foraging. Let us consider a simple case, in which the amount of daylight *d* can just take two values of 0 and 1 (corresponding to night and day), and there are just two types of trees which give a payoffs *p* of 0 or 1. Let us further assume that approaching a tree has a cost of *n* = 0.2. Figure 8A shows how the reward in this task depends on the amount of daylight and fruit density: the overall reward is equal to 1 − *n* only for approaching a fruit-rich tree during daytime, and *−n* otherwise. Let us consider a task in which on each trial an animal is presented with an opportunity to forage on a tree and needs to decide whether to approach it or not.

**Figure 8:**
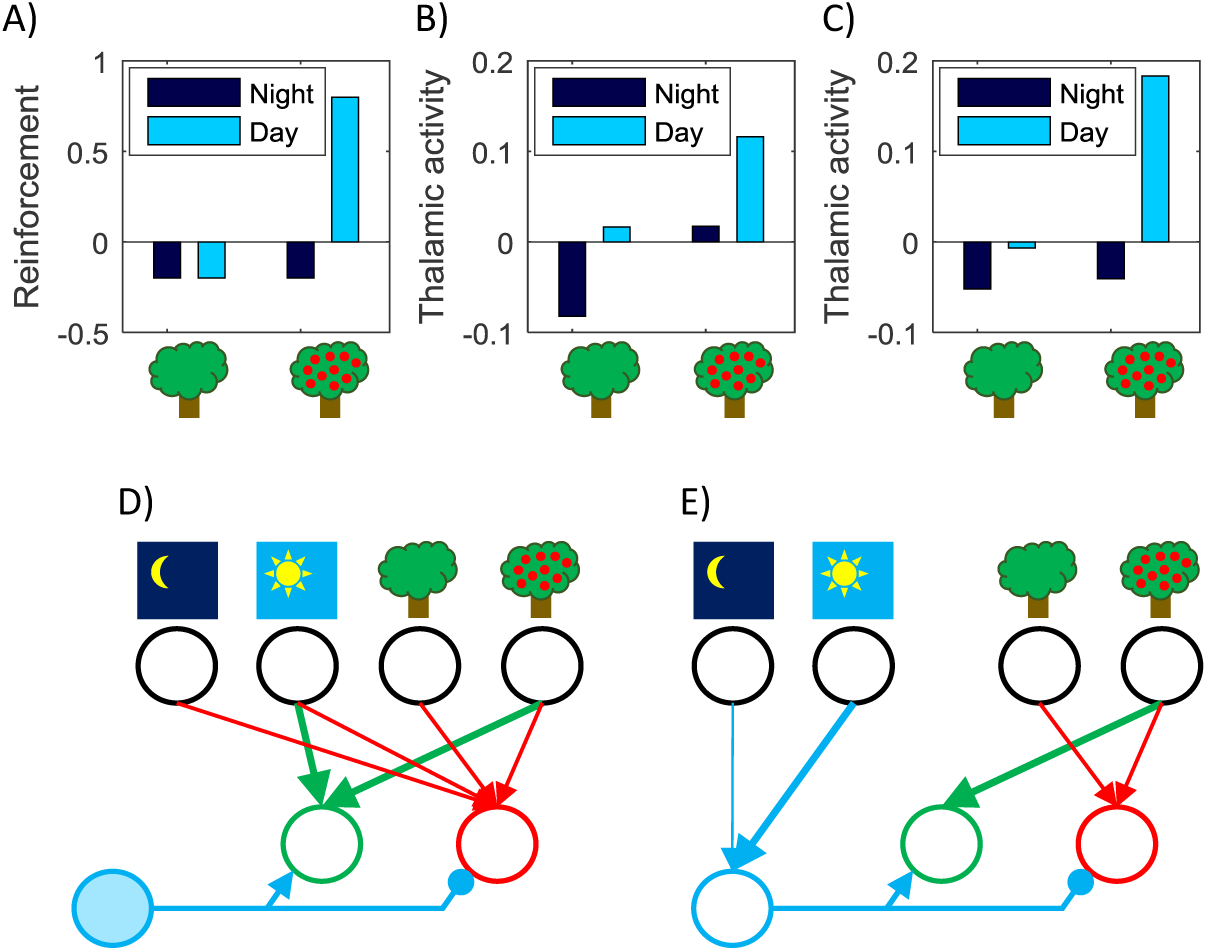
Learning the foraging policy depending on the amount of daylight. A) Total reinforcement for approaching trees with different fruit density at different times of day. B, C) Average thalamic activity in the models which did not (B) and did (C) learn the dopaminergic motivation signal. The standard errors of thalamic activity were smaller than 0.01, so the error bars are not shown. D, E) Architectures of the model that did not (D) and did (E) learn the dopaminergic motivation signal. Black circles denote sensory populations, blue circles denote dopaminergic neurons, and green and red circles denote the Go and No-Go neurons. The thickness of the arrows reflect the average values of weights learned by each model. For the model in panel D these weights were equal to: *G* = [0, 0.19, 0, 0.19] and *N* = [0.09, 0.07, 0.09, 0.07]. For the model in panel E these weights were equal to: *G* = [0, 0.23], *N* = [0.06, 0.07] and *w* = [0.1, 0.84].

Before showing how this task can be effectively solved, let us first analyse the difficulties faced by a model in which the dopaminergic motivation signal is independent from the state of the environment. Figure 8D shows the architecture of this model. It includes four populations providing information on the state of the environment, so on each trial two of these populations were active. All sensory neurons send projections to the Go and No-Go neurons, and the thickness of connections indicates the values of synaptic weights that have been learned by the model (see Materials and Methods). Positive values of Go weights have been learned only from sensory units selective for daytime and high fruit density, because payoff was only received when they were active. All No-Go weights converged to similar values, as effort was independent from the state of the environment. In this model the dopaminergic neuron did not received input from sensory populations so the dopaminergic motivation signal was equal to a constant baseline value. Figure 8B shows the thalamic activity produced by the model for different states of the environment. It was lowest for the fruitless tree in the night as then the Go neurons did not receive any input, and highest for the fruit-rich tree during daytime, when the Go neurons received input from two sensory populations (Figure 8D). In the other two states (fruit-rich tree in the night and fruit-less tree during daytime), the Go neurons received input from one sensory population, so the thalamic activity in these intermediate states was the average of the states with the lowest and highest activity. Thus the model produced positive thalamic activity in the intermediate states, although approaching tree in these states gave overall negative reinforcement. This simulation illustrates that a linear model may be unable to learn the optimal policy determined by a non-linear utility function of Equation 15.

Figure 8E shows a network architecture in which the sensory populations encoding the amount of daylight project to the dopaminergic rather than striatal neurons. Here again the thickness of arrows indicates the strength of learned connections, and shows that dopaminergic neuron learned to produce higher activity during day than night. Figure 8C shows the thalamic activity produced by the model. It is positive only for approaching fruit-rich tree during daytime. The model does not approach it in the night, because low dopaminergic motivation signal leads to under-weighting payoffs and over-weighting costs. Thus thanks to the dependence of dopaminergic motivation signal on daylight, the model was able to learn the optimal policy.

Dopaminergic neurons have also been observed to produce increased activity in the proximity of a reward (Howe et al., 2013). Figure 9A re-plots the dopamine concentration observed in a sample trial in a T-maze task (Howe et al., 2013), which gradually increases closer to the reward. Such response of dopaminergic neurons may arise because, when the reward is close, there is a higher probability of obtaining it, so executing actions is more useful in general, and it is beneficial to put the basal ganglia in an energised state.

**Figure 9:**
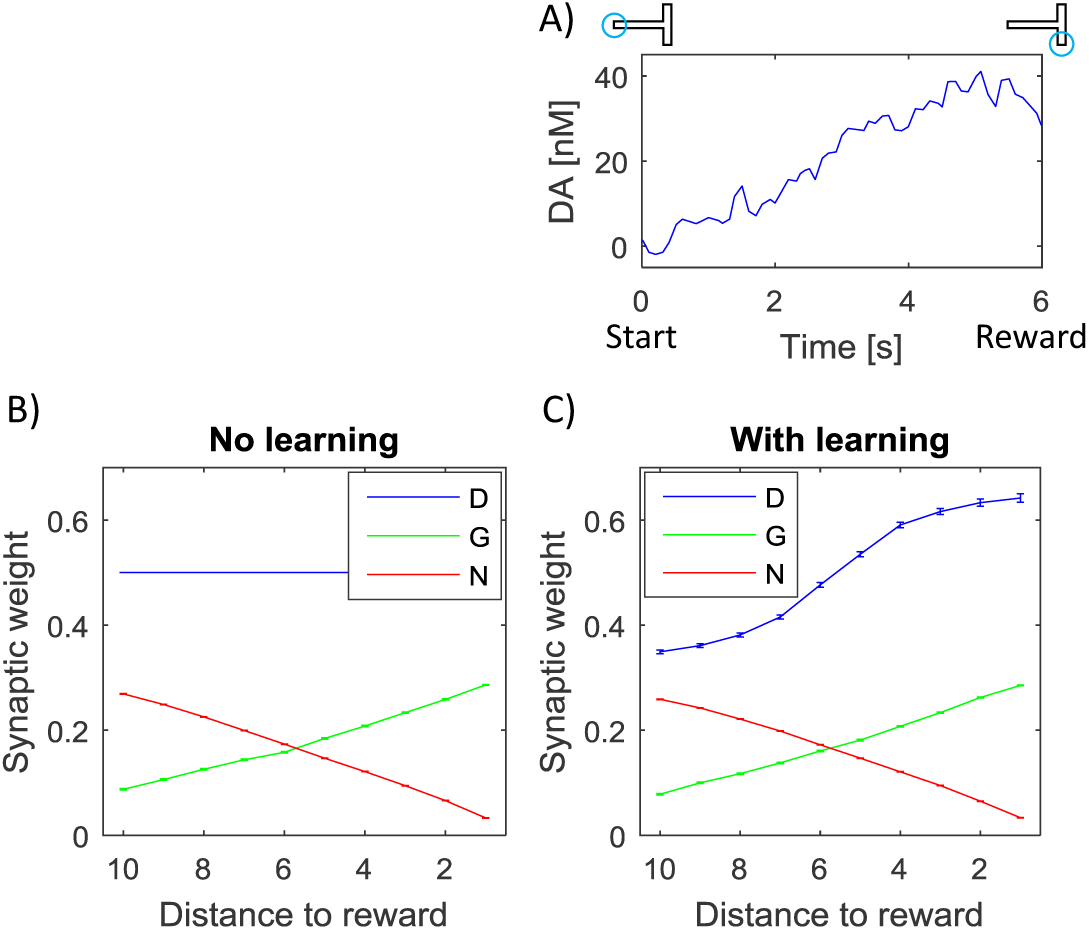
Learning the dopaminergic motivation signal at different proximities to reward. A) Concentration of dopamine measured in striatum during a sample trial in a T-maze. At time 0 the Go cue is presented, while at time 6 the animal reaches the reward. Data re-plotted from Figure 1b of paper by Howe et al. (2013). B, C) Simulations of learning. In each panel, the green and red curves indicate the weights of Go and No-Go neurons from the neurons selective for a particular distance to the reward. The blue lines show the weights *w* of dopaminergic neurons which determine the average dopaminergic motivation signal *D*. The curves show the average weight values at the end of 1000 simulations, and the error bars show the standard error. B) Simulations without learning of dopaminergic motivation signal. C) Simulations with learning.

To see more clearly the benefit of increasing dopaminergic motivation signal in proximity of reward, let us consider a task in which at the beginning of each trial an animal is a certain number of steps away from a reward with payoff *p*, and needs to decide whether to approach it or not. Approaching each step has a cost *n*. Furthermore, during each time step the reward can disappear with certain probability (which could correspond to being eaten by another animal). Thus in this task, the utility of approaching the fruit is equal to:

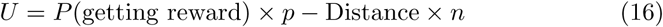

The two scaling factors in the above equation depend on the proximity in the opposite ways. Namely, the probability of getting the reward increases with proximity, while the Distance is just the opposite of proximity. Thus the above equation has the same structure as the thalamic activity defined in Equation 2, so this utility could be evaluated by the basal ganglia if dopaminergic neurons were modulated by the reward proximity (see Section 1 of Supplemental Information). Analogously as for the case of daylight, Figure S1 shows an example of a task where the optimal policy cannot be learned by a linear model with fixed dopaminergic motivation signal, but can be found when the dopaminergic neurons learn how their motivation signal should depend on reward proximity.

We demonstrated that there exists a category of tasks in which learning the dopaminergic motivation signal is necessary for finding the optimal policy. However, when faced with a new task, dopaminergic neurons “do not know a priori” if the task belongs to this category. Thus the dopaminergic neurons may learn to adjust its motivation signal on the basis of the state of environment even if it not necessary for finding the optimal policy (as long as such adjustment does not impair reward). To illustrate this property, let us consider a special case of the above task in which all fruits give the same payoff, as then the optimal policy is simply determined by the distance to the reward. For example, with *p* = 1, *n* = 0.1 and the probability of reward disappearing equal to 0.1, the optimal policy is to approach for distances smaller than 6. The optimal behaviour in this task can be learned by the reinforcement learning model described in the previous subsection, without any additional adjustments in the motivation signal, and Figure 9B shows the resulting striatal weights when the dopaminergic motivation signal was fixed to the baseline value. The weights of Go neurons are higher than the weights of No-Go neurons for distances smaller than 6, so the model produces the optimal policy. When the dopaminergic neuron is allowed to learn, it does so to produce a higher motivation signal closer to reward. This is shown in Figure 9C and resembles the experimentally observed increase in dopamine in reward proximity (Howe et al., 2013). This additional learning does not have any decremental affect, as the model still produces the optimal policy.

To understand the reasons behind the observed dopaminergic responses in various experimental studies, it is insightful to analyse how the level of dopaminergic motivation signal learned with the Reinforce model depends on the relative value of acting. A simple scenario was simulated in which an animal was in a particular state where only one action was available. If the animal performed the action, it incurred a cost (e.g. due to effort) of *n* = 0.5, and received a subsequent reinforcement *r*_*act*_. If the animal did not perform the action, it received reinforcement *r*_*no act*_. We considered combinations of *r*_*act*_ and *r*_*no act*_ from a range [ −1, 1]. For each combination, we simulated an animal learning a single set of weights of Go (*G*), No-Go (*N*), and dopaminergic neurons determining the average motivation signal (*D*). The resulting weights are shown in Figure 10. The difference in the striatal weights converges to a value proportional to the total reinforcement for selecting an action, i.e. *r*_*act*_ *− n*. By contrast, the dopaminergic motivation signal converges to the values above baseline when the overall reward for executing action is higher than doing nothing i.e. *r*_*act*_ *− n > r*_*no act*_, and to the values below baseline otherwise.

**Figure 10:**
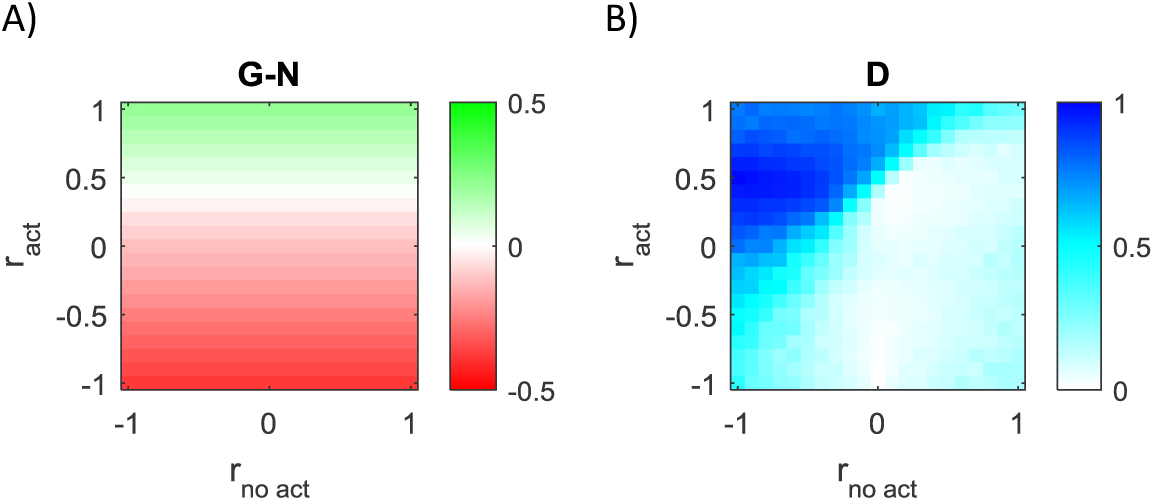
Synaptic weights learned when the reinforcements were provided for acting and not acting. In each panel the co-ordinates correspond to reinforcement for acting *r*_*act*_ and for not acting *r*_*no act*_. The colour visualizes the average weight values at the end of simulations. A) The difference between weights *G* of Go neurons and *N* of No-Go neurons. B) The weights *w* of the dopaminergic neuron that determine the average dopaminergic motivation signal *D*.

Figure 10B illustrates that a high level of dopaminergic motivation signal can be produced in situations in which expected overall reward is negative, e.g. when one may need to take an action to avoid an aversive outcome (*r*_*act*_ = 0, *r*_*no act*_ = *-*1). Although the value of acting is equal to zero, it is higher than value of not acting, so the simulated dopaminergic neuron learned to produce motivation signal above baseline. An analogous increased level of dopamine in the striatum has been observed after cues predicting aversive stimuli, that could be avoided by pressing a lever, on trials when a rat avoided them (Oleson et al., 2012). Conversely, a low dopaminergic motivation signal can be learned in a situation with high expected reward, e.g. in a No-Go paradigm in which an animal needs to refrain from making any movements in order to obtain a reward (*r*_*act*_ = 0, *r*_*no act*_ = 1). As making an action is undesirable in this task, the simulated animal learns to have the dopaminergic motivation signal below baseline. An analogous low level of dopamine has been experimentally measured following a cue after which the animal was required to remain still for 2s in order to obtain a reward (Syed et al., 2016). A low motivation signal has also been learned in Figure 10B in a case when an aversive outcome was delivered irrespective of the animal’s actions (*r*_*act*_ = −1, *r*_*no act*_ = −1). In this situation the total reinforcement for acting *r*_*act*_ -*n* includes the cost of acting, so it is even lower than the total reinforcement for not acting *r*_*no act*_. An analogous low level of dopamine has been observed in the striatum after a cue that had been paired with an unavoidable shock (Oleson et al., 2012).

The simulations in Figures 9 and 10 demonstrate that once the dopaminergic neurons are allowed to learn, they start to produce responses depending on the benefit of moving, reminiscent of several experimentally observed patterns of responses. The responses of dopaminergic neurons observed in other situations could be accounted for in an analogous way. For example, the responses to salient stimuli (Redgrave and Gurney, 2006) could arise from the fact the salient stimuli often require a reaction, e.g. saccade to find out what is going on. Similarly, responses to novel stimuli (Schultz, 1998) or reward uncertainty (Fiorillo et al., 2003) may be connected with the fact that in such situations it is useful to explore to find out about consequences of actions, so that appropriate actions may be taken in the future (Kakade and Dayan, 2002). The average dopamine level has also been shown to correlate with the rate of receiving rewards (Hamid et al., 2016). In a situation where many rewards are available, it is useful to move to gather rewards quickly (Niv et al., 2005), so a high dopaminergic motivation signal could be learned.

## Discussion

This article proposed how the basal ganglia select actions on the basis of past experience and the current motivational state. First, it suggests that the basal ganglia computes a utility function in which the positive and negative consequences encoded in striatal neurons are differentially weighted by the dopaminergic motivation signal. Second, it describes how the positive and negative consequences of actions can be separately learned if the dopaminergic neurons also encode reward prediction error. Third, it suggests that dopaminergic neurons themselves learn what level of activity they need to produce to optimize behaviour. In this section we relate the theory with experimental data and other models, state experimental predictions, and highlight the directions in which the theory needs to be developed further.

### Information encoded by dopaminergic neurons

The proposed theory assumes that firing of dopaminergic neurons encodes information about two quantities: motivation *m* and reward prediction error *δ*. The motivation signal is determined by long-lasting factors such as hunger, but on top of this it is also influenced by the necessity to make movements in the current state, which may change on a fast time scale during behaviour. Encoding of both motivation and prediction error by dopaminergic neurons is consistent with a study of Syed et al. (2016). In this experiment in order to obtain reward, the rats needed to press a lever after certain cues, and remain still after different cues. The study was designed such that the need to move and the reward prediction error were de-correlated across conditions. The dopamine concentration in the striatum during the task could only be explained by a combination of movement and reward prediction error signals (Syed et al., 2016).

Another recent study (Hamid et al., 2016) reported that on a time-scale of seconds, the dopamine concentration not only encodes the reward prediction error but is also strongly correlated with the time-discounted value of expected reward (i.e. increases with temporal reward proximity). Hamid et al. (2016) stated in the title of their paper that “dopamine signals the value of work”. In the simulations shown in the present paper the motivation signal *D* was also correlated with the value of an action, encoded in the difference in striatal weights *G − N*. In Figure 9C the motivational signal *D* (blue curve) is positively correlated with *G* (green) and negatively correlated with *N* (red). There also exists a positive correlation between *D* and *G − N* across simulated conditions in Figure 10 as high motivation signal was mainly learned in a region where the value of action was higher. Thus the proposed theory agrees with Hamid et al. (2016) that the dopaminergic motivation signal is correlated with the value of executing actions, but here it is proposed that the motivation signal also depends on other factors.

Given that dopamine release may encode both motivation and learning signals on fast time-scales, a question arises of how the striatal neurons distinguish which signal is being transmitted at the moment and whether they should change their activity or synaptic connections. As mentioned in the Result section, other neuromodulators such as acetylcholine may provide information on when the learning needs to take place. Such a role of acetylcholine is consistent with the observation that the striatal cholinergic interneurons pause when feedback is provided, irrespective of the outcome (Morris et al., 2004). Furthermore, the reduced concentration of acetylcholine has been proposed to be necessary for the plasticity of striatal neurons (Nair et al., 2015). However, cholinergic neurons are likely to also be involved in generating the motivation signal, as the antagonists of cholinergic receptors are used for treatment of Parkinson’s disease, suggesting that they increase the overall tendency to select actions (Katzenschlager et al., 2002). Furthermore, the cholinergic neurons seems to have a complex effect on dopamine release (Rice and Cragg, 2004). Nevertheless, if two neuromodulators are both involved in modulating striatal activity and plasticity, it is conceivable that in the two dimensional space of their concentrations both motivation and teaching signals are encoded, and the details of this encoding may be clarified by future studies.

### Relationship to other theories

Given the success of the reinforcement learning theory in predicting dopaminergic responses in learning tasks, several researchers have asked if all responses of dopaminergic neurons occurring on a fast time scale could encode reward prediction error. For example, it has been demonstrated that after making additional plausible assumptions the reinforcement learning models could produce a prediction error signal with a dynamics resembling the dopaminergic activity observed after salient and novel stimuli (Kakade and Dayan, 2002), and in reward proximity (Kato and Morita, 2016). Furthermore, Kato and Morita (2016) have shown that reducing prediction error in their model produces similar effects as dopamine depletion in tasks studied by Salamone et al. (2016). However, certain patterns of dopaminergic activity are challenging to account for by the prediction error alone. For example, in a Go-No-Go task, dopamine release differed between conditions with the same reward and time to reward depending on whether the movement was necessary (Syed et al., 2016). Also, increased dopamine release was observed after stimuli predicting avoidable aversive outcome, and this increase started before the rat pressed the lever to avoid the shock (Oleson et al., 2012). The stimuli signalled transition from a neutral state to a state with a negative expected value, so the reward prediction error was negative, and yet the dopamine concentration increased. As mentioned earlier, these patterns of activity are naturally explained by assuming that dopamine also encodes a motivation signal depending on a relative value of making movements.

Recently, there has been a debate concerning the fundamental concept of basal ganglia function, i.e. the relationship between the Go and No-Go neurons: on one hand they have the opposite effects on a tendency to make movements (Kravitz et al., 2010), but on the other hand they are co-activated during action selection (Cui et al., 2013). The presented theory is consistent with both observations: It assumes that Go and No-Go neurons have opposite effects on movement initiation. But during action selection the basal ganglia need to calculate the utility which combines information encoded by both populations, so may require their co-activation.

The proposed model builds on the seminal work of Collins and Frank (2014), who proposed that the Go and No-Go neurons learn the tendency to execute and inhibit movements, and how the level of dopamine changes the influence of the Go and No-Go pathways on choice. The key new feature of the present model is the ability to learn both payoffs and costs associated with an action. It has been illustrated in Figure 5 that when the model repeatedly selects an action resulting first in a cost and then in the payoff, *G* and *N* converge to values proportional to *p* and *n*. In the OpAL model, the Go and No-Go neurons can learn the payoff and cost if an action has only payoffs or only costs. Figure S2 shows that when the OpAL model repeatedly selects an action resulting first in a cost and then in the payoff, Go and No-Go weights converge to zero (this is also shown analytically in Section 6 of Supplemental Information).

To simulate the effects of dopamine depletion on choice between an arm of a T-maze with more pellets behind a barrier and an arm with with fewer pellets, Collins and Frank (2014) trained a model on three separate actions: eating in the left arm, eating in the right arm, and crossing a barrier. In this way it was ensured that each action had just payoff or just cost, and the model could learn them. Subsequently, during choice the model was deciding between a combination of two actions (e.g. crossing a barrier and eating in the left arm) and the other action. By contrast, the model proposed in this paper was choosing just between the two options available to an animal in an analogous task (Figure 2), because it was able to learn both payoffs and costs associated with each option. This is a useful ability, as most real world actions have both payoffs and costs.

As the direction of weight changes in the proposed model is the same as in the OpAL model, it retains the ability to describe some of the phenomena explained by the OpAL model. For example, Beeler et al. (2012) trained rats to stay on a rotating rod. Control animals were able to learn the task, while after blocking dopaminergic receptors they were not. Interestingly, after washing out of the antagonists, the animals took longer to learn the task than naive animals (Beeler et al., 2012). Collins and Frank (2014) reproduced this effect in simulations, in which during training with a dopamine blocker, the simulated animals received negative feedback (as they were unable to perform this task due to motor difficulties) resulting in an increase in No-Go weights and a decrease in Go weights. Thus during subsequent training without the blocker, it took longer for the weights of the Go neurons corresponding to the correct action to increase to the level allowing selecting it sufficiently quickly. Figure S3 shows that the proposed model shows analogous dynamics of weights as OpAL in such simulations.

In the original paper introducing the plasticity rules (Mikhael and Bogacz, 2016), it was proposed that the rules allow the Go and No-Go neurons to encode reward variability, because when an action results in variable rewards, both *G* and *N* increase during learning. It was further proposed that the tonic level of dopamine controls the tendency to make risky choices, as observed in experiments (Rutledge et al., 2015), because it leads to emphasizing potential gains, and under-weighting potential losses. However, here it is proposed that the striatal learning rules and dopaminergic motivation signal primarily sub-serve a function more fundamental for survival, i.e. learning payoffs and costs and biasing how they affect thalamic output in order to compute the utility function of Equation 1. From this perspective, the influence of dopamine level on tendency to make risky choices arises as a by-product of a system primarily optimized to choose actions that maximize utility.

### Experimental predictions

Each part of the theory described in the three subsections of the Results makes testable predictions. First, similarly as the OpAL model (Collins and Frank, 2014), the theory proposes that the positive and negative consequences are separately encoded by the Go and No-Go neurons which are differentially modulated by dopamine. The theory predicts that agonists specific to just one of the striatal populations (e.g. a D2 agonist), should decrease the effect of consequences encoded by this population (e.g. negative) without changing the impact of the other population. This prediction could be tested in an experiment involving choice between options with both payoff and cost. In particular, the theory predicts that the degree of preference of a neutral option (*p* = 1, *n* = 1) over a high cost option (*p* = 1, *n* = 2) should increase with D2-agonist, while the preference of a high payoff option (*p* = 2, *n* = 1) over a neutral option (*p* = 1, *n* = 1) should not be affected by the D2-agonist.

It would be interesting to investigate whether changing the influence of positive and negative consequences on choice can not only be achieved by pharmacological manipulations, but also by changing a behavioural context such as hunger, or reward rate which has been shown to affect the average dopamine level (Hamid et al., 2016). If such an experiment was done in humans (or non-human primates), an eye-tracker could be used to investigate whether participants spend more time on a part of the stimulus informing about payoff in blocks with high hunger or reward rate.

Second, the theory assumes that the synaptic plasticity rules include a decay term proportional to the value of the synaptic weights themselves. Decay terms are also present in other models of learning in basal ganglia (Franklin and Frank, 2015; Yttri and Dudman, 2016; Kato and Morita, 2016). This class of models predicts that the synaptic weights of striatal neurons which are already high increase less during potentiation than the smaller weights (an opposite prediction is made by the OpAL model (Collins and Frank, 2014), where the weights scale the prediction error in the update rule). This prediction could be tested by observing the Excitatory Post-Synaptic Currents (EPSCs) evoked at individual spines. The class of model including decay predicts that the spines with smaller evoked EPSCs before inducing plasticity should be more likely to potentiate.

Third, the theory proposes that the dopaminergic neurons learn what level of activity they need to produce. If they employ a learning algorithm similar to Reinforce, then one should be able to observe a relationship between the activity of dopaminergic neurons on consecutive trials analogous to those illustrated in Figure 7. For example, if average activity of dopaminergic neurons was relatively high on trial *t* and the reward was better than expected, then the activity on trial *t* + 1 should be higher than that expected from random fluctuations.

### Extensions of the theory

There are multiple directions in which the presented theory could be extended. For example, the theory has to be integrated with the models of action selection in the basal ganglia to describe how the circuit selects the action with the highest utility. It also needs to be demonstrated how the theory accounts for the symptoms of the disorders of the basal ganglia. It has to be described how the utility is computed in the part of basal ganglia involving ventral striatum which has a slightly different organization (Kupchik et al., 2015). The definition of utility can be extended to incorporate the cost of waiting to allow the model to maximize the reward rate (Niv et al., 2005). Furthermore, the theory may be extended to describe the dependence of the dopaminergic teaching signal on the motivational state (Cone et al., 2016).

While defining utility in Equation 1, we assumed for simplicity thatthe utility depends linearly on payoffs and cost. However, according to the prospect theory (Kahneman and Tversky, 1979), payoffs and costs are each transformed by a different non-linear function before being combined into the utility. The responses of dopaminergic neurons to conditioned stimuli, also reflect the non-linear utility function rather than the expected reward (Stauffer et al., 2014).

Furthermore, in the utility defined in Equation 1 the motivation has a single dimension. But there are multiple dimensions of motivation and corresponding consequences, e.g. hunger and food, thirst and water, etc. (Keramati and Gutkin, 2014). Complex animals like humans additionally have more abstract motivational drives that also include multiple dimensions (Maslow, 1943). While defining the utility, we assumed for simplicity that the motivation only affects positive consequences, but certain aspects of negative consequences are affected by their corresponding motivation dimension, e.g. the effort associated with an action affects utility differentially depending on the level of tiredness.

Taking the above arguments into account, the definition of utility could be extended:

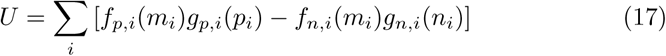

In the above equation *m*_*i*_, *p*_*i*_, *n*_*i*_ denote dimension *i* of motivation and consequences, *f*_*p,i*_ and *f*_*n,i*_ are the functions determining how the dimension *i* of motivation scales the positive and negative consequences (see Section 1 of Supplemental Information), and *g*_*p,i*_ and *g*_*n,i*_ are the non-linear functions, that determine the subjective value of payoffs and costs in the prospect theory.

Let us consider how plausible it is for the basal ganglia to evaluate this extended utility. First, different dimensions of motivation would have to be encoded by different groups of dopaminergic neurons or neurons releasing different neuromodulators. Second, the striatal neurons receiving *m*_*i*_ would have to encode the corresponding *p*_*i*_ and *n*_*i*_. This could arise if the same groups of neurons providing motivational signal *m*_*i*_ also provided the teaching signal *δ*_*i*_ allowing the striatal synapses to learn about consequences in the corresponding dimension. For example, the noradrenergic neurons may provide both *m*_*i*_ and *δ*_*i*_ signals connected with the dimension of effort. The norepinephrine level increases with tiredness (Hartley et al., 1972), and the phasic changes in firing rate of noradrenergic neurons encode the information about the effort associated with currently performed action (Varazzani et al., 2015).

It is intriguing to ask if the evaluation of utility of actions combining separately encoded positive and negative consequences is also performed by areas beyond the basal ganglia. Indeed, positive and negative associations are encoded by different populations of neurons in the amygdala (Namburi et al., 2015). Furthermore different cortical regions preferentially project to Go or No-Go neurons (Wall et al., 2013), raising the possibility that the positive and negative consequences are also encoded separately in the cortex. It would be interesting to investigate to what extent the extended utility describes the actual computations in the brain, and if so what dimensions (or their combinations) are encoded by different neurons and neuromodulators.

## Materials and Methods

### Finding the values to which the striatal weights converge

Here we calculate to what values the striatal synaptic weights converge when updated according to Equations 13 and 14. When an action has a cost *n* and payoff *p*, the weights are updated twice: with *r* = *−n* after making an effort, and *r* = *p* after the payoff. Thus the weight changes are approximately equal to:

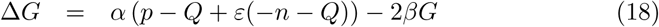

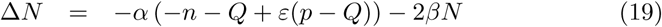

In the above equations *Q* denotes the expected reward, i.e. *Q* = *G − N*. The above equations are approximate, because after the first update (with *r* = *−n*), *Q* also changes which is not considered here for simplicity, but note that this change becomes small as the learning and decay rates *α* and *β* decrease.

As *Q* appears in update rules of Equations 18 and 19, let us first analyse to what value it converges. Subtracting Equations 18 and 19 we obtain:

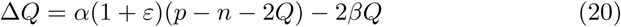

At convergence, *Q* no longer changes, thus setting Δ*Q* = 0 and solving the above equation for *Q* we obtain its value at the fixed point:

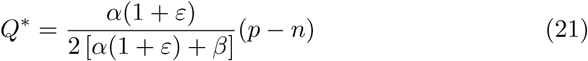

We see that *Q* converges to a value proportional to *p − n*. Denoting the proportionality constant by *c*_1_, and substituting to Equation 18 we obtain:

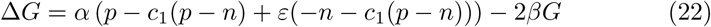

After convergence *G* does not change, so setting Δ*G* = 0 and solving for *G* we find its value at the fixed point:

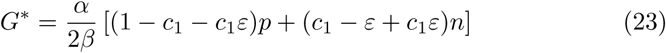

In order for *G* to be proportional just to *p*, the term scaling *n* must be equal to 0, i.e.

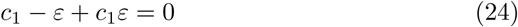

Substituting the definition of *c*_1_ and rearranging terms, we obtain the condition the parameters of the learning rules need to satisfy for *G*^*∗*^ *∼ p*:

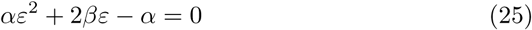

It is interesting to note that When *α* = *β*, there is a unique value of *ε* satisfying the above condition 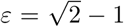 ≈ 0.41.

By symmetry, when the condition of Equation 25 is satisfied, then *N* ^*∗*^ *∼ n*. In summary, when the parameters satisfy Equation 25, then the weights of Go and No-Go neurons converge to the values proportional to *p* and *n* respectively.

It is also interesting to note that when Equation 24 is satisfied, then Equation 23 becomes:

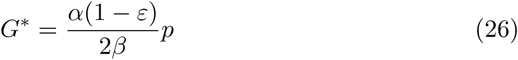

Analogously, *N* ^*∗*^ would be proportional to *n* with the same proportionality constant. The model converges to such fixed points in the limit of small *α* and *β*, while for higher constants the value of *G* tends to be higher and *N* tends to be slightly lower; see Figure 5, where the positions of the fixed points are indicated by dashed lines. This discrepancy comes from the approximation made in Equations 18 and 19 that the expected reward during the second update (with *r* = *p*) is also *Q* (see the paragraph under Equations 18 and 19). By contrast, the expected reward during the second update is lower because it was decreased during the first update (with *r* = *−n*). Lower expected reward results in higher prediction error and higher *G*.

### Simulations of the effects of dopamine level on choices

During simulations of an experiment by Salamone et al. (1991), the model received payoff *p*_1_ = 10 for choosing a pellet, and payoff *p*_2_ for approaching the lab chow. The model was simulated in two conditions differing in the cost of choosing a pellet which was equal to *n*_1_ = 0 in the free-pellet condition, and to *n*_1_ = *n*_*lever*_ in a condition requiring lever pressing to obtain a pellet. There was no cost of choosing lab chow *n*_2_ = 0. For each condition, the model was simulated in two dopamine states: in the intact state the dopaminergic motivation signal was equal to a baseline value during choice *D* = 0.5 while in the state corresponding to the presence of dopamine antagonist it was set to a lower value *D* = *D*_*anta*_.

For each condition and state, the behaviour of *N*_*rats*_ was simulated. Each simulation consisted of *N* training and *N* testing trials, where *N* = 180 (as each animal in the experiment of Salamone et al. (1991) was tested for 30 minutes, so 180 trials corresponds to an assumption that a single trial took 10s). At the start of each simulation the weights were initialized to *G*_*i*_ = *N*_*i*_ = 0.1. During each training trial the model experienced choosing a pellet (i.e. received cost *n*_1_, modified weights *G*_1_ and *N*_1_, and then received payoff *p*_1_ and modified the weight again), and approaching the lab chow. During each testing trial, the thalamic activity for each option was calculated from Equation 2, and Gaussian noise with standard deviation *σ* was added. An option with the highest thalamic activity was selected, and if this activity was positive, the action was executed, resulting in the corresponding cost and payoff and weight modification. If thalamic activity for both options was negative, no action was executed and no weights were updated.

The values of model parameters: *p*_2_, *n*_*lever*_, *D*_*anta*_, *σ* were optimized to match the choices made by the animals. In particular, for each set of parameters, the model was simulated *N*_*rats*_ = 100 times, and the average number of choices 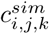 of option *i* in dopamine state *j* and experimental condition *k* was computed. The mismatch with corresponding consumption in experiment 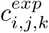 was quantified by a normalized summed squared error:

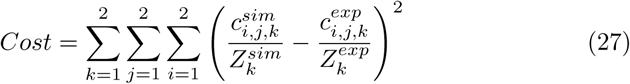

In the above equation 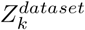 is a normalization term equal to the total number of choices or consumption in a particular condition:

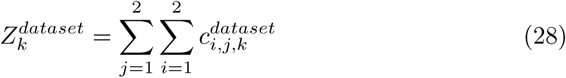

The values of parameters minimizing the cost function were sought using the Simplex optimization algorithm implemented in Matlab, and the following values were found *p*_2_ = 2.34, *n*_*lever*_ = 7.11, *D*_*anta*_ = 0.37, *σ* = 0.38. Subsequently, the model with these optimized parameters was simulated with *N*_*rats*_ = 6, which was the number of animals tested by Salamone et al. (1991). The resulting mean number of choices and the standard error across animals are shown in Figure 6.

### Reinforce model

According to the Reinforce model (Williams, 1992), on a trial where the dopaminergic neuron produced activity *D*, the weights *w* from neurons encoding the current state are modified on the basis of how the reinforcement *r* differed from the estimated mean reinforcement *V* in the current state:

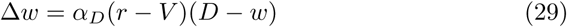

In the above equation *α*_*D*_ is a learning rate constant. The term (*r − V*) in the above equation would correspond to the reward prediction error in the actor-critic framework (see Section 4 of Supplemental Information). Hence in that framework the plasticity rule is local in the sense that all variables occurring in it are available to a dopaminergic neuron, as it can access the information from its teaching signal, motivation signal and synaptic weights. In the main text, for simplicity we do not consider the critic, and we do not simulate how *V* is learned. It has been demonstrated that the Reinforce algorithm works well even if the average reward *V* is misestimated (Williams, 1992), so for simplicity we set it to 0, and use a simplified version of the rule:

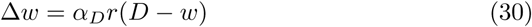

### Simulations of the effects of daylight

Each simulation consisted of 1000 trials, and the simulations were repeated 100 times. At the start of each simulation the striatal weights were initialized to 0. The models illustrated in Figures 8D and E included four sensory nodes *x*_*i*_ which were set to 1 in states corresponding to night, day, fruitless tree and fruit-rich tree, respectively.

In the model in Figure 8D all sensory nodes sent projections to Go and No-Go neurons with weights *G*_*i*_ and *N*_*i*_, and dopaminergic motivation signals was set to a baseline value *D* = 0.5. On each simulated trial the animal had to decide whether to approach a tree. Thus the activity of thalamus was computed:

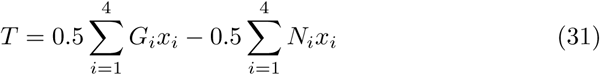

Additionally a Gaussian noise with mean 0 and standard deviation 0.1 was added to the above thalamic activity. If the thalamic activity was positive, the tree was approached, resulting in initially negative reinforcement *r* = *−*0.2 due to cost and subsequent reinforcement that was *r* = 1 if a fruit-rich tree was approached during daytime or *r* = 0 otherwise. After each reinforcement, the striatal weights originating from active sensory popu*v*lations were modified according to Equations 13 and 14 with *α* = *β* = 0.05, 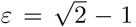, and prediction error equal to:

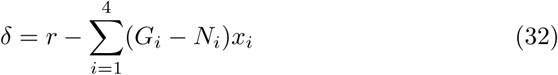

In the model in Figure 8E the sensory nodes selective for the amount of daylight projected to dopaminergic neurons with weights *w*_*d*_, which were initialized to 0.5 at the start of each simulation. The sensory nodes selective for the amount of fruits sent projections to Go and No-Go neurons with weights *G*_*i*_ and *N*_*i*_ respectively. At the start of each trial with daylight *d*, the dopaminergic motivation signal was set to *D* = *w*_*d*_ and additional Gaussian noise with mean 0 and standard deviation 0.2 was added. The thalamic activity was determined on the basis of a pair of weights *G*_*i*_ and *N*_*i*_ from sensory population selective for the fruit density *i* on a given trial, according to Equation 2. As in the case of the model described earlier in this subsection, Gaussian noise with standard deviation 0.1 was added to the thalamic activity, and if the thalamic activity was positive, the tree was approached, resulting in the pattern of reinforcements described above.

After each reinforcement, the striatal weights originating from the active sensory population were modified according to Equations 13 and 14 with the same parameters as before, and prediction error based on these weights, defined according to Equation 12. Additionally, on each trial, irrespectively whether the action was taken or not, the weight *w*_*d*_ of dopaminergic neuron was modified according to Equation 30 with *α*_*D*_ = 0.2 and *r* taken as the total reinforcement in that trial. Throughout the simulations all weights were constrained to nonnegative values, and dopaminergic weights were additionally constrained not to exceed 1.

### Simulations of the effects of reward proximity

Each simulation consisted of 1000 trials. On each simulated trial, the animal was located at a certain distance *d* ϵ {1, *…,* 10*}* from the reward. Over the course of simulations the animal was learning weights *G*_*d*_, *N*_*d*_ of striatal neurons, and *w*_*d*_ of dopaminergic neurons describing the strength of connections from neurons selective for being at a distance *d* from the reward. At the beginning of each simulation the weights were initialized to *G*_*d*_ = *N*_*d*_ = 0, and *w*_*d*_ = 0.5.

On each simulated trial the animal had to decide whether to approach the reward. The dopaminergic motivation signal *D* was set to *w*_*d*_, and noise with normal distribution with mean 0 and standard deviation 0.1 was added to it. Next the thalamic activity was set according to Equation 2, and noise with mean 0 and standard deviation 0.1 was also added to allow exploration. If *T >* 0, the animal approached the reward, otherwise it did not.

If the animal approached the reward, it incurred cost *n* = 0.1*d* and could receive payoff *p* = 1 with probability 0.9^*d*^, corresponding to the assumption of probability 0.1 of reward disappearing while traversing a unit of distance. The weights of striatal neurons were thus modified according to Equations 13 and 14 twice (with *r* = *−n* and *r* = *p*) with parameters *α* = *β* = 0.05 and 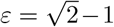. If the animal decided not to approach the reward, then no cost or payoff was given.

In the simulations in Figure 9B, after each trial the weights of the dopaminergic neurons were modified according to Equation 30 based on total reward *r* = *p − n*, with learning rate *α*_*D*_ = 0.4.

### Simulations with reinforcement for acting and not acting

These simulations were performed analogously to the one described in the previous subsection. At the beginning of each simulation the weights were initialized to *G* = *N* = 0, and *w* = 0.5. The thalamic activity was generated in the same way as described in the previous subsection, and an action was selected when *T > −*0.1. A negative value of the threshold was used to allow execution of actions to avoid a negative outcome. A negative threshold on *T* can be neurally implemented by adding a positive constant input to thalamic activity (defined in Equation 2) and executing actions when such modified thalamic activity is positive.

If the animal took the action, it incurred cost *n* = 0.5 and subsequently received reinforcement *r*_*act*_. Thus the weights of striatal neurons were modified according to Equations 13 and 14 twice (with *r* = *−n* and *r* = *r*_*act*_) with parameters *α* = *β* = 0.1 and *ε* = 2 *−* 1. If the animal decided not to approach the reward, then it received reinforcement *r*_*no act*_, but the striatal weights were not modified.

After each trial the weights of the dopaminergic neurons were modified according to Equation 30 based on the total reward (*r* = *r*_*act*_ *− n* if action was taken and *r* = *r*_*no act*_ otherwise), with learning rate *α*_*D*_ = 0.4. If as a result of the modification, the weight *w* became negative, it was set to *w* = 0, and if *w* exceeded 1, it was set to *w* = 1. For each combination of *r*_*act*_ and *r*_*no act*_, 100 simulations were performed with 1000 trials in each. The average values of the weights at the end of the simulation are reported in Figure 10.

## Acknowledgements

This work was supported by the Medical Research Council grant MC UU 12024/5. The author thanks Paul Dodson, John Mikhael and Mark Walton for reading an earlier version of the manuscript, and Peter Brown, Masud Hus-sain, Joshua Berke, Sanjay Manohar, Stephanie Cragg, Chris Summerfield, Alex Kacelnik, Kevin Lloyd, Jean Debarros, Alek Pogosyan, Tim Behrens, and Peter Magill for discussion.

